# Skeletal muscle myosin heavy chain protein fragmentation as a potential marker of protein degradation in response to resistance training and disuse atrophy

**DOI:** 10.1101/2024.05.24.595789

**Authors:** Daniel L. Plotkin, Madison L. Mattingly, Derick A. Anglin, J. Max Michel, Joshua S. Godwin, Mason C. McIntosh, João G. A. Bergamasco, Maíra C. Scarpelli, Vitor Angleri, Lemuel W. Taylor, Darryn S. Willoughby, C. Brooks Mobley, Andreas N. Kavazis, Carlos Ugrinowitsch, Cleiton A. Libardi, Michael D. Roberts

**Author notes:** address co-correspondence to: Michael D. Roberts, PhD, Auburn University Endowed Alumni Professor, Director, Nutrabolt Applied and Molecular Physiology Laboratory, School of Kinesiology, Auburn University, 301 Wire Road, Office 286, Auburn, AL 36849, Cleiton A. Libardi, PhD, MUSCULAB - Laboratory of Neuromuscular Adaptations to Resistance Training/Department of Physical Education/Federal University of São Carlos – UFSCar., Rod. Washington Luiz, km 235 – SP 310, CEP 13565-905, São Carlos, SP, Brazil. indicates D.L.P. and M.D.R. primarily drafted manuscript.

## Abstract

We sought to examine how resistance exercise (RE), cycling exercise, and disuse atrophy affect myosin heavy chain (MyHC) protein fragmentation in humans. In the first study (1boutRE), younger adult men (n=8; 5±2 years of RE experience) performed a lower body RE bout with vastus lateralis (VL) biopsies obtained immediately before, 3-, and 6-hours post-exercise. In the second study (10weekRT), VL biopsies were obtained in untrained younger adults (n=36, 18 men and 18 women) before and 24 hours (24h) after their first/naïve RE bout. These participants also engaged in 10 weeks (24 sessions) of resistance training and donated VL biopsies before and 24h after their last RE bout. VL biopsies were also examined from a third acute cycling study (n=7) and a fourth study involving two weeks of leg immobilization (n=20, 15 men and 5 women) to determine how MyHC fragmentation was affected. In the 1boutRE study, the fragmentation of all MyHC isoforms (MyHC_Total_) increased 3 hours post-RE (∼ +200%, p=0.018) and returned to pre-exercise levels by 6 hours post-RE. Immunoprecipitation of MyHC_Total_ revealed ubiquitination levels remained unaffected at the 3- and 6-hour post-RE time points. Interestingly, a greater increase in magnitude for MyHC type IIa versus I isoform fragmentation occurred 3-hours post-RE (8.6±6.3-fold versus 2.1±0.7-fold, p=0.018). In all 10weekRT participants, the first/naïve and last RE bouts increased MyHC_Total_ fragmentation 24h post-RE (+65% and +36%, respectively; p<0.001); however, the last RE bout response was attenuated compared to the first bout (p=0.045). The first/naïve bout response was significantly elevated in females only (p<0.001), albeit females also demonstrated a last bout attenuation response (p=0.002). Although an acute cycling bout did not alter MyHC_Total_ fragmentation, ∼8% VL atrophy with two weeks of leg immobilization led to robust MyHC_Total_ fragmentation (+108%, p<0.001), and no sex-based differences were observed. In summary, RE and disuse atrophy increase MyHC protein fragmentation. A dampened response with 10 weeks of resistance training, and more refined responses in well-trained men, suggest this is an adaptive process. Given the null polyubiquitination IP findings, more research is needed to determine how MyHC fragments are processed. Moreover, further research is needed to determine how aging and disease-associated muscle atrophy affect these outcomes, and whether MyHC fragmentation is a viable surrogate for muscle protein turnover rates.

## INTRODUCTION

Our laboratory recently recruited college-aged men with prior resistance training experience to perform two lower body resistance exercise (RE) bouts separated by one week consisting of 30% versus 80% one repetition loads [1]; these being termed 30-FAIL and 80-FAIL bouts, respectively. Vastus lateralis (VL) muscle biopsies were obtained immediately prior to as well as 3- and 6-hours following these bouts, and we sought to holistically examine skeletal muscle-molecular outcomes that differed between the two loading paradigms. Our first series of experiments indicated that both bouts similarly altered global DNA methylation and transcriptome-wide markers [1]. Our second report indicated that both bouts similarly increased certain aspects of the mechanistic target of rapamycin signaling complex 1 (mTORC1) cascade while also similarly increasing follistatin mRNA and protein expression [2]. Notably, both studies support prior literature suggesting that low-load and high-load training elicit similar post-exercise anabolic signaling outcomes so long as sets are performed near failure [3-5].

The final phase of project analysis began with utilizing the remaining 30-FAIL tissue for 8 participants to examine if titin phosphorylation was altered 3- or 6-hours following exercise. Our interest was spawned by past reviews suggesting this phenomenon may be a catalyst for post-exercise anabolic signaling [6, 7]. To accomplish this aim, myofibrils were isolated and solubilized using our recently published MIST method adopted by us and others [8-11]. During pilot SDS-PAGE Coomassie experiments with myofibril isolates, we observed notable protein fragmentation occurred 3 hours following exercise that was visibly reversed by the 6-hour post-exercise time point (depicted in Figure 1 in the Results section). With preliminary immunoblotting experiments we also observed a similar trend with the titin protein, thus precluding phosphorylation analysis. After a thorough examination of the literature, only one paper has reported that a RE bout promotes titin and myosin heavy chain (MyHC) fragmentation 3 hours following a RE bout [12], and this report was in seven previously trained men. However, aside from briefly mentioning this as being potential evidence of protein disruption with resistance training, the significance of this finding was not further explored.

**Figure 1.**
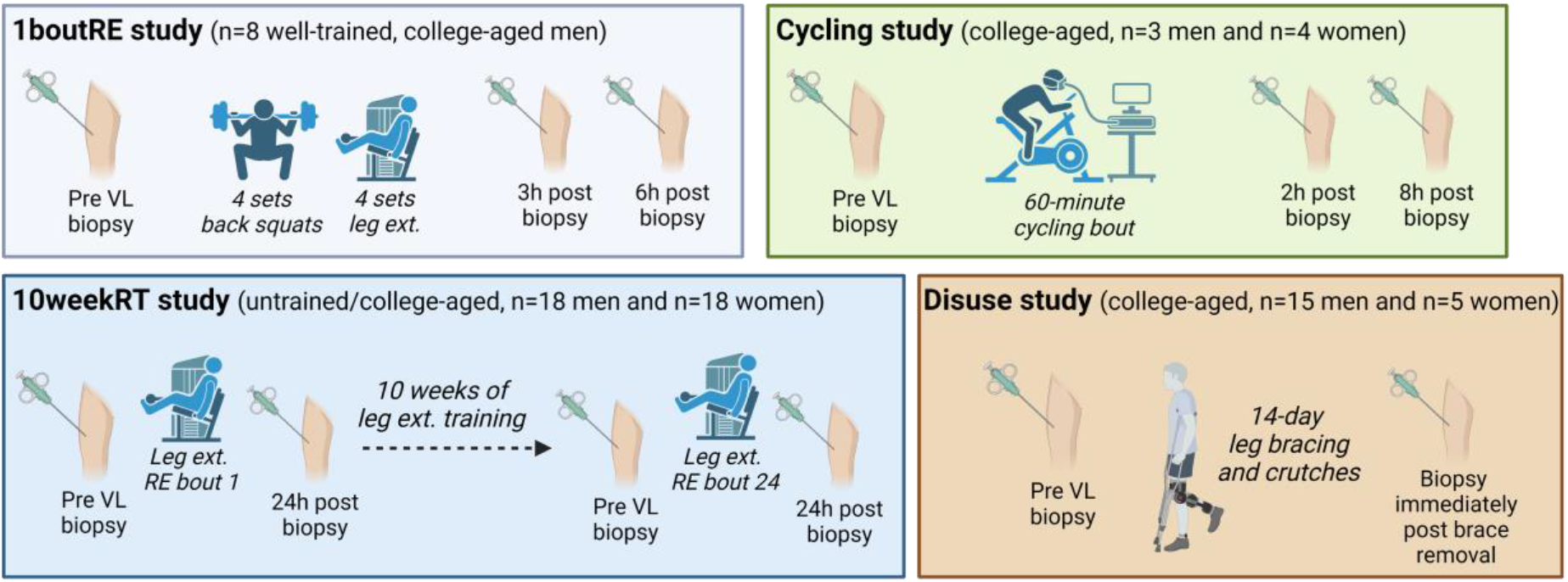
Summary of human studies Schematic (drawn using Biorender.com) illustrates the study logistics and participant number for each study whereby MyHC analyses occurred. More details related to each study can be found in-text.

These observations motivated a series of exploratory experiments using VL biopsy specimens from various human studies. Data from one study (termed “1boutRE”) provides compelling evidence that significant MyHC fragmentation occurs 3 hours following a single lower body RE bout in well-trained males. However, MyHC fragments are largely absent 6 hours following exercise implying that skeletal muscle can rapidly clear these proteins following a loading stimulus. In a second study (termed “10weekRT”), we observed that MyHC fragmentation is present 24 hours following a leg extensor bout in a large cohort of untrained males and females, and that this 24-hour response is attenuated after 10 weeks (24 total sessions) of leg extensor resistance training. Results from our third study indicated that 60 minutes of cycling exercise did not promote MyHC fragmentation 2- or 8-hours post-exercise. Results from our fourth study indicated that two weeks of disuse atrophy through leg immobilization promoted a robust MyHC fragmentation response. We believe that this easy-to-perform immunoblot-based technique could be used as a proxy marker of protein degradation in resistance exercise studies or disuse studies. Experimental details and an expanded discussion of these findings are provided in the following paragraphs.

## METHODS

### 1boutRE study participants

Muscle specimens from well-trained college-aged males (n=8; 22±3 years old, 5±2 years of RE experience, 83.0±7.0 kg, 1.6±0.3 1RM squat: body mass) as described by Sexton et al. [1], and all experimental procedures were approved by the Auburn University Institutional Review Board (IRB protocol #20-081). Information regarding participant characteristics, the acute lower body RE exercise bout, and the procurement of vastus lateralis biopsies can be found in Sexton et al. [1]. Briefly, participants reported to the laboratory during morning hours in a fasted state. After donating a baseline biopsy, participants performed 4 sets each of the back squat and leg extension exercises at 30% of their estimated one-repetition maximum loads until volitional failure. Five minutes of rest was allowed between sets and exercises. Following the RE bout, VL biopsies were collected 3- and 6-h post-exercise. This study, along with others detailed herein, are visually depicted in Figure 1.

### 10weekRT study participants

Muscle specimens were from healthy college-aged participants (n=38 total with 19 women [24.2 ± 4.9 years old, 62.7 ± 8.5 kg and 1.64 ± 0.1 m) and 19 men (24.5 ± 3.3 years old, 73.6 ± 13.4 kg, 1.76 ± 0.1 m) as previously described by Scarpelli et al. [13]. Due to sample limitations for 2 participants, only 36 participants were analyzed. All experimental procedures were approved by the local ethics committee, the study was conducted in accordance with the most recent version of the Declaration of Helsinki and was registered as a clinical trial (Brazilian Registry of Clinical Trials – RBR-57v9mrb), and training as well as specimen collection was performed at the University of Sao Carlos. The resistance training protocol consisted of four sets of 9–12 maximum repetitions of unilateral leg extension exercises, with a 90-second rest period between sets. The load was adjusted for each set to ensure that concentric muscle failure occurred within the target repetition range. Participants completed 24 training sessions over a period of 10 weeks, with sessions conducted 2 to 3 times per week. Critically, four mid-thigh VL biopsies were obtained before (Pre) and 24 h after the first training bout (untrained state), and 96 hours after the second to last training bout (Pre) and 24 h after the last training bout (trained state).

### Cycling study and leg immobilization study participants

To determine how an acute cycling bout affects post-exercise MyHC fragmentation, human muscle specimens from a previously published study from our laboratory were analyzed (IRB protocol #18-226) [14]. To determine how non-complicated (i.e., without injury or illness) disuse atrophy affects MyHC fragmentation, human muscle specimens from another ongoing Auburn University IRB-approved study (IRB protocol #23-220) were analyzed. Experimental procedures from both studies were approved by the Auburn University Institutional Review Board and were conducted at Auburn University.

For the cycling study, apparently healthy college-aged participants (n=7, 3 males and 4 females; 23±3 years old, 23.0±2.9 kg/m^2^) reported to the laboratory during the morning hours under fasted conditions and donated a baseline VL biopsy. Participants then mounted a cycle ergometer (Velotron, RacerMate, Seattle, WA, USA) and performed a 5-minute warm-up at a self-selected pace. Wattage was adjusted thereafter to achieve 70% VO_2_ reserve and participants cycled for 60 minutes. Post-exercise biopsies were then obtained 2- and 8-hours following the cycling bout.

For the leg immobilization study, apparently healthy college-aged participants (n=20, 15 males and 5 females; 26±3 years old, 25.9±5.6 kg/m^2^) reported to the laboratory under fasted conditions and donated a baseline VL biopsy. Participants were then fitted with a knee brace locked at 90º and administered crutches and explicit instructions to prevent weight-bearing activities on the braced leg for a 14-day period. Following the 14-day disuse period, participants reported back to the laboratory under fasted conditions (± 2 hours from the first visit) and donated a second VL biopsy.

### 1boutRE tissue myofibril and cytoplasmic fractionation

Using a liquid nitrogen-cooled ceramic stage, approximately 20 mg of muscle from each biopsy specimen was placed in 1.7 mL tubes containing 300 μL of ice-cold homogenizing buffer (Buffer 1: 20 mM Tris-HCl, pH 7.2, 5 mM EGTA, 100 mM KCl, 1% Triton-X100; all chemicals from VWR; Radnor, PA, USA). Samples were homogenized using tight-fitting pestles and centrifuged at 3,000 g for 30 minutes at 4ºC. Supernatants (cytoplasmic fraction) were transferred to new 1.7 mL tubes and stored at -80ºC until protein analyses described below. As a wash step, resultant pellets (myofibrillar fraction) were resuspended in Buffer 1, and samples were centrifuged at 3,000 g for 10 minutes at 4ºC. Resultant supernatants from this step were discarded, myofibril pellets were resuspended in 300 μL of ice-cold wash buffer (Buffer 2: 20 mM Tris-HCl, pH 7.2, 100 mM KCl, 1 mM DTT; all chemicals from VWR), and tubes were centrifuged at 3,000 g for 10 minutes at 4ºC; this step was performed twice. Final myofibril pellets were resuspended in 400 μL of ice-cold storage buffer (Buffer 3: 20 mM Tris-HCl, pH 7.2, 100 mM KCl, 20% glycerol, 1 mM DTT, 50 mM spermidine; all chemicals from VWR), and stored at -80°C for analyses described below.

### Whole tissue lysate preparations for 10weekRT, cycling, and leg immobilization studies

Using a liquid nitrogen-cooled ceramic stage, approximately 20 mg of muscle from each biopsy specimen was placed in 1.7 mL tubes containing 400 μL of commercially available general cell lysis buffer (Cell Signaling; Danvers, MA, USA; cat#: 9803). Samples were centrifuged at 500 g for 5 minutes at 4ºC. Resultant supernatants from placed in new 1.7 mL tubes and stored at - 80°C for analyses described below.

### MyHC immunoblotting

Protein concentrations of 1boutRE myofibril and cytoplasmic isolates, and whole tissue lysates from the other studies were quantified using bicinchoninic acid (BCA) colorimetric assays (Thermo Scientific, Waltham, MA, USA). Isolates and lysates from all studies were then prepared for Western blot analysis with 4× Laemmli buffer for final concentration preparations at 1 μg/μL. Aliquots of prepared samples (4 μL for myofibrillar preps, 15 μL for cytoplasmic preps, and 10 μL of whole tissue lysate preps) were applied to 4−15% SDS-polyacrylamide gels (Bio-Rad; Hercules, CA, USA) and subjected to electrophoresis at 180 volts for 50 minutes in a preformulated 1× SDS-PAGE buffer (VWR). Proteins were then electrotransferred onto pre-activated polyvinylidene difluoride membranes (Bio-Rad) for 2 hours on ice, Ponceau stained, and placed in a gel documentation system (ChemiDoc Touch; Bio-Rad) to capture whole-lane images for protein normalization purposes. Membranes were then blocked in a solution containing 5% skimmed milk powder in Tris-buffered saline with 0.1% Tween-20 (VWR) for 1 hour at ambient temperature.

Incubation of membranes with anti-MyHC antibodies were carried out for 1-2 hours (room temperature) at a dilution of 1:200 in TBST containing 5% bovine serum albumin. These antibodies included non-concentrated clone supernatants of: i) mouse monoclonal IgG2a MyHC, termed MyHC_Total_ throughout (Developmental Studies Hybridoma Bank; Iowa City, IA, USA; cat#: A4.1025), ii) mouse monoclonal IgG1 MyHCI (Developmental Studies Hybridoma Bank; cat#: A4.951), iii) mouse monoclonal IgG1 MyHCIIa (Developmental Studies Hybridoma Bank; cat#: SC-71), and iv) mouse monoclonal IgM MyHCIIx (Developmental Studies Hybridoma Bank; cat#: 6H1). Following primary antibody incubations, membranes were washed for 15 minutes in TBST and incubated with horseradish peroxidase-conjugated anti-mouse IgG (Cell Signaling; cat#: 7076) or IgM (ThermoFisher Scientific; cat#: 31440) secondary antibodies at a dilution of 1:2000 in TBST with 5% BSA for one hour (room temperature) prior to development steps described below.

Membranes were developed for 1-5 seconds using an enhanced chemiluminescence reagent (Luminata Forte HRP substrate; Millipore Sigma) in a gel documentation system (ChemiDoc Touch; Bio-Rad). The densitometry of prominent MyHC bands and fragments were quantified with ImageLab v6.0.1 (Bio-Rad) using the “Lanes & Bands Tool” functions. Densitometry readings for bands and targets were normalized to baseline (Pre) values, which were averaged to a value of 1.00, and expressed as fold-change from Pre.

### Immunoprecipitation for MyHC_Total_ polyubiquitination in 1boutRE myofibril isolates

To determine if MyHC fragments were polyubiquitinated, immunoprecipitation (IP) experiments were performed on 1boutRE myofibril isolates using a commercially available kit (Dynabeads Protein G; Thermo Fisher Scientific; cat#: 10009D). Per sample reaction, 50 μL of resuspended bead slurry was mixed with 30 μL of mouse monoclonal IgG2a MyHC (Developmental Studies Hybridoma Bank; cat#: supernatant of A4.1025) for 60 minutes at room temperature on an inversion apparatus. Bead-IgG2a complexes were washed with antibody binding/wash buffer provided by the kit and subsequently incubated with 600 μg of myofibril protein per sample for 60 minutes at room temperature on an inversion apparatus. Bead-Ab-Ag complexes were then washed three times with wash buffer provided by the kit, and 20 μL of elution buffer as well as 10 μL of 4× Laemmli buffer was added. Samples were boiled for 5 minutes at 100ºC, beads were removed using a magnetic rack apparatus, and immunoblotting experiments were carried out on 10 μL of resultant IP preps whereby polyubiquitinated MyHC fragments were probed using a polyclonal rabbit IgG antibody (1:1000; Cell Signaling; cat#: 3933). In addition to these IP experiments, 1boutRE myofibril isolates were immunoblotted for polyubiquitination using the same polyclonal rabbit IgG antibody and immunoblotting methods described in the prior section.

### Statistical analysis

Stats were performed and graphs were constructed using commercially available software (GraphPad Prism, v10.1.0; Boston, MA, USA). Most 1boutRE and all cycling study data were analyzed via one-way repeated measures ANOVAs. When significant model effects were observed (p<0.05), Tukey’s post hoc tests were performed to determine which time points were significantly different from one another. The only 1boutRE data analyzed via two-way repeated measures ANOVAs were isoform-specific data. When significant model effects were observed (p<0.05), Fisher’s LSD post hoc tests were performed to determine which time points were significantly different from one another. All 10weekRT data were analyzed via two-way (training status × time) repeated measures ANOVAs. When significant model effects were observed (p<0.05), Tukey’s post hoc tests were performed to determine which time points were significantly different from one another. Leg immobilization study data were analyzed using dependent samples t-tests. Data throughout are presented as means and standard deviation values with individual data points.

## RESULTS

### Evidence of post-exercise myofibril protein fragmentation following a single RE bout

As noted in the Introduction section, our experiments began with attempting to interrogate the titin protein from 1boutRE specimens. Figure 2 shows preliminary SDS-PAGE Coomassie experiments on myofibril isolates in two participants from this study. Notably, visual protein fragments were observed in the MyHC region 3 hours following the RE bout, and the rapid disappearance of these fragments was evident by 6 hours post-exercise.

**Figure 2.**
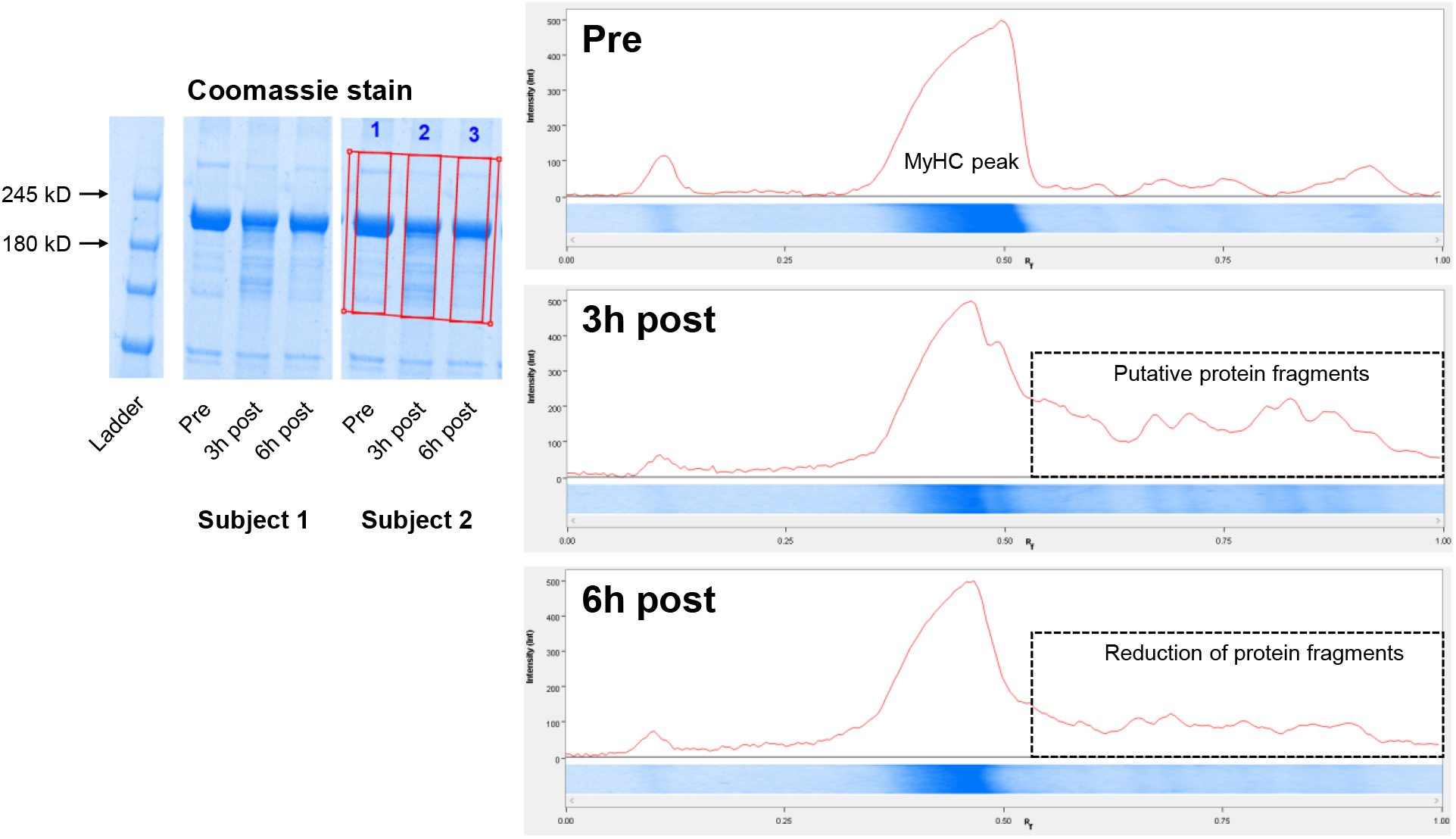
Evidence of post-exercise myofibril protein fragmentation in 1boutRE study participants As discussed in-text, preliminary 1boutRE experiments were performed on two well-trained participants’ myofibril isolates aiming to examine the presence of titin using 4-15% SDS-PAGE gels and Coomassie staining. In both participants, visual myofibril protein fragments were observed in the myosin heavy chain (MyHC) kilodalton region 3 hours following the resistance exercise bout. Conversely, the rapid disappearance of these fragments was evident by the 6-hour post-exercise time point.

### Transient post-exercise MyHC_Total_ fragmentation following a single RE bout

Figure 3 shows MyHC_Total_ immunoblotting experiments in all 1boutRE participants. The increased presence of MyHC_Total_ fragmentation was evident in the myofibril fraction 3 hours following RE, and the rapid disappearance of these fragments was evident by the 6-hour post-exercise time point (Figure 3a). MyHC_Total_ fragmentation was also observed at the 3-hour post-exercise time point in the cytoplasmic fraction of several participants (Figure 3b), albeit this did not reach statistical significance.

**Figure 3.**
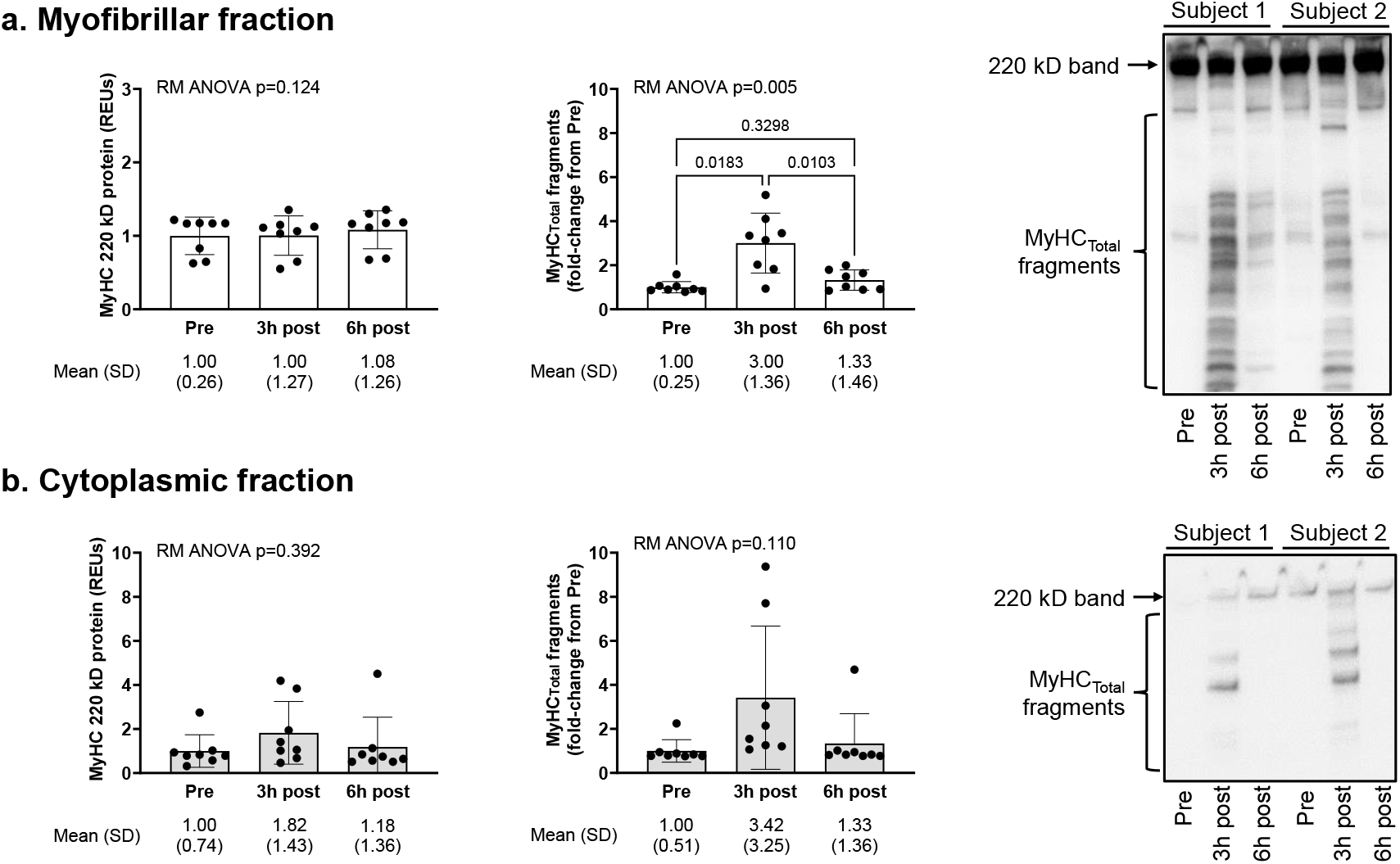
Post-exercise MyHC_Total_ fragmentation in 1boutRE participants Data from well-trained 1boutRE men (n=8) show that significant total myosin heavy chain (MyHC_Total_) fragmentation is evident in the myofibril fraction 3 hours following a resistance exercise bout (panel a); however, the rapid (and significant) disappearance of these fragments was evident by the 6-hour post-exercise time point. Also notable is the high presence of MyHC_Total_ fragments in the cytoplasmic fraction in several participants (panel b); however, this did not reach statistical significance. Representative immunoblots are shown for 2 of 8 participants, and data are presented as mean and standard deviation values with repeated measures (RM) ANOVA p-values.

### Isoform-specific MyHC fragmentation following a single RE bout

Figure 4 shows isoform specific MyHC immunoblotting experiments in all 1boutRE participants. The increased presence of type I and IIa MyHC fragments were evident in the myofibril fraction 3 hours following the RE bout, and the rapid disappearance of these fragments was evident by the 6-hour post-exercise time point (Figure 4a). Additionally, the magnitude of 3-hour post-RE type IIa isoform fragmentation was greater than type I isoform fragmentation (p=0.024). Finally, there were visually different patterns of fragmentation between isoforms, with lighter molecular weight type I isoform fragments appearing post-RE versus heavier type IIa fragments (Figure 4b).

**Figure 4.**
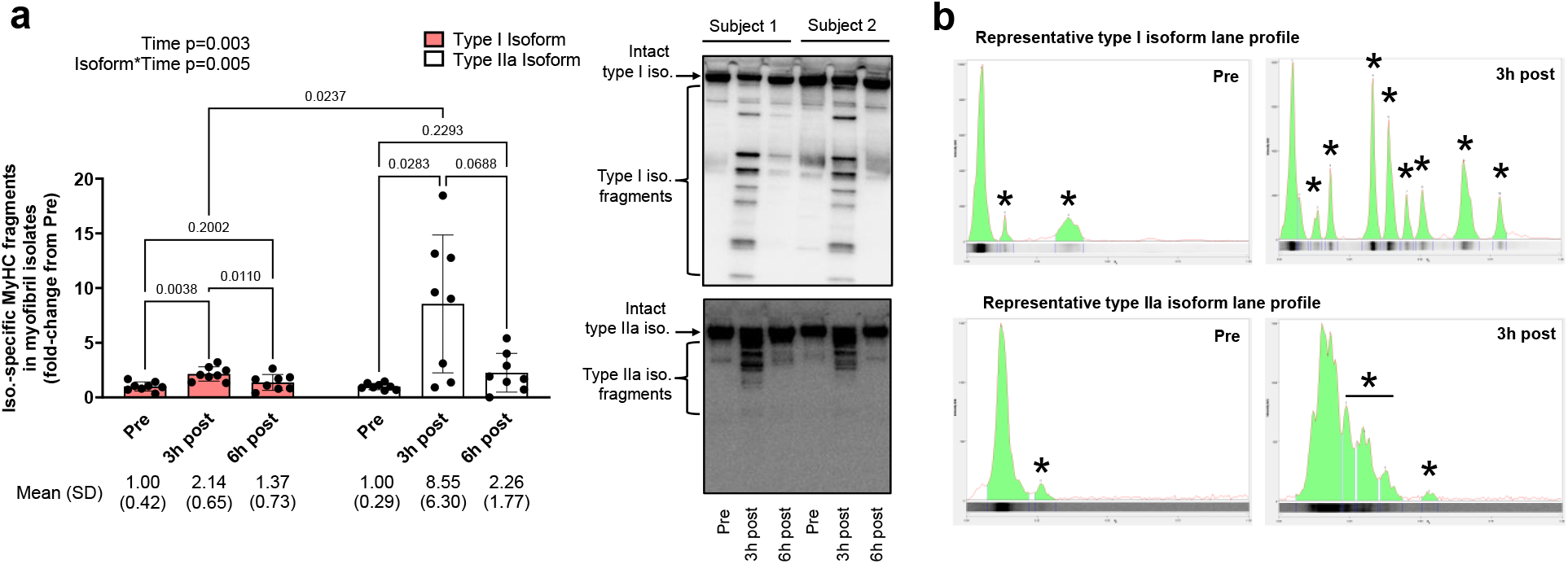
Type I versus IIa MyHC isoform fragmentation in 1boutRE participants Data from well-trained 1boutRE men (n=8) show that significant myosin heavy chain (MyHC) fragmentation of the type I and IIa isoforms is evident in the myofibril fraction 3 hours following the resistance exercise bout (panel a); however, as with MyHC_Total_ fragments, the rapid (and significant) disappearance of I and IIa fragments was evident by the 6-hour post-exercise time point. Also notable were the different patterns of fragmentation between isoforms, with lighter molecular weight type I isoform fragments appearing post-RE versus heavier type IIa fragments. Representative immunoblots are shown for 2 of 8 participants, and data are presented as mean and standard deviation values with two-way (isoform*time) ANOVA time and interaction p-values. Panel b shows lane profiles of type I and IIa isoform fragmentation from two different participants where “*” indicates fragments detected by analysis software.

### Polyubiquitination of myofibril proteins and MyHC_Total_ poly-ubiquitination following a single RE bout

Figure 5 shows myofibril protein polyubiquitination and MyHC_Total_ polyubiquitination in all 1boutRE participants. Myofibril protein poly-ubiquitination levels did not significantly differ between pre- and post-exercise timepoints (Figure 5a). Moreover, polyubiquitinated fragments in the ∼15-50 kD region (where the signal was prominent) were not significantly altered when this signal was normalized to the MyHC IP signal (Figure 5b). Also notable is the lack of polyubiquitinated MyH_Total_ fragments between the ∼50-220 kD region.

**Figure 5.**
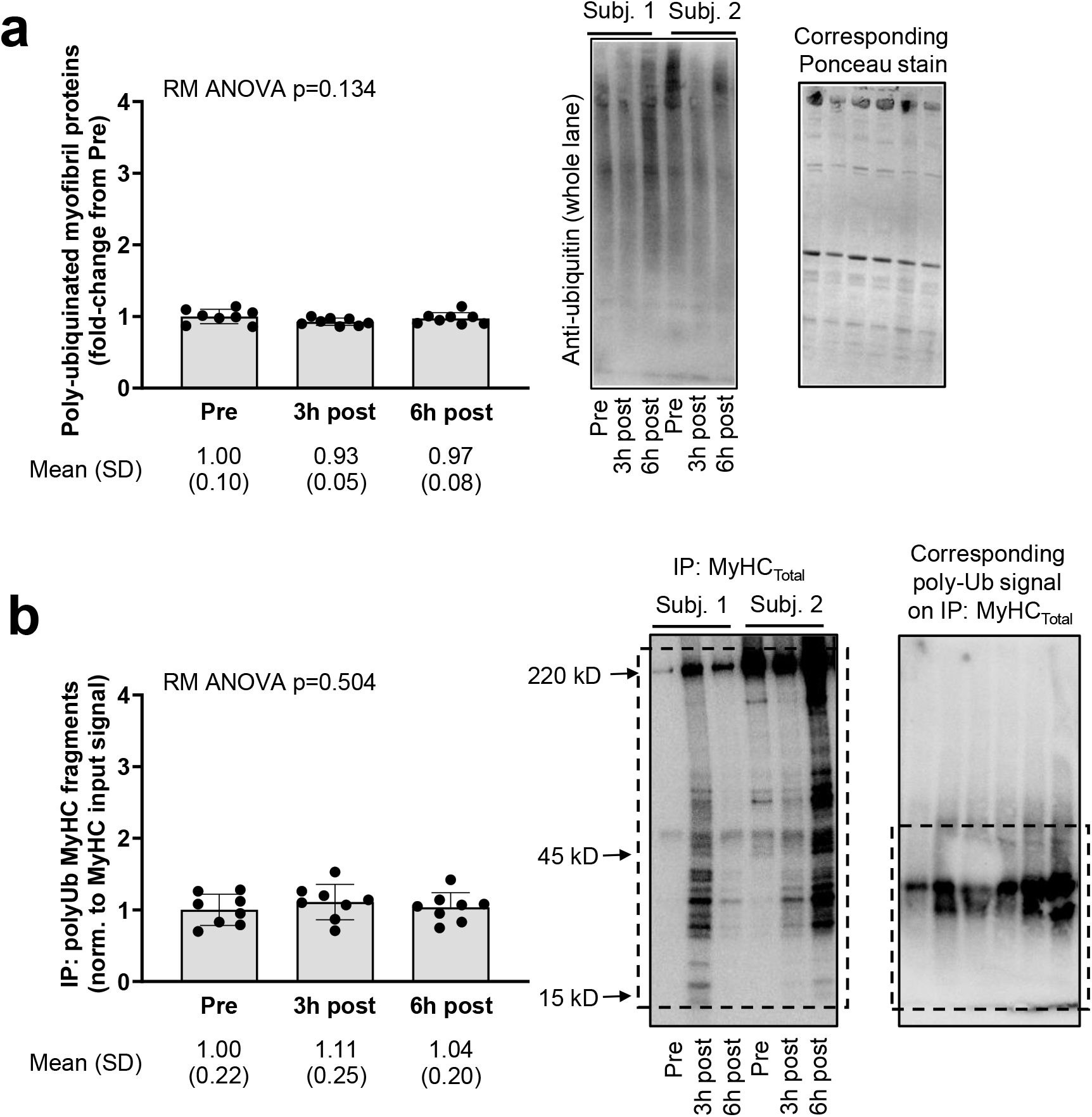
Total myofibril protein and MyHC_Total_ poly-ubiquitination in 1boutRE participants Data from well-trained 1boutRE men (n=8) show that total myofibril protein polyubiquitination levels remain unaltered post-exercise (panel a). Additionally, the polyubiquitination signal on immunoprecipitated MyHC fragments (spanning ∼15-50 kD) remained unaltered 3- and 6-hours post-exercise when data were normalized to the IP: MyHC signal (panel b). Representative immunoblots are shown for 2 of 8 participants, and data are presented as mean and standard deviation values with one-way repeated measures (RM) ANOVA p-values.

### Post-exercise MyHC_Total_ fragmentation in 10weekRT participants in the naïve and trained states

Figure 6 shows MyHC_Total_ fragmentation responses in 10weekRT participants. Significant increases were observed 24-hours following the first/naïve and last training bouts (Figure 6a). However, this response was attenuated following the last versus the first/naïve bout (p=0.045).

**Figure 6.**
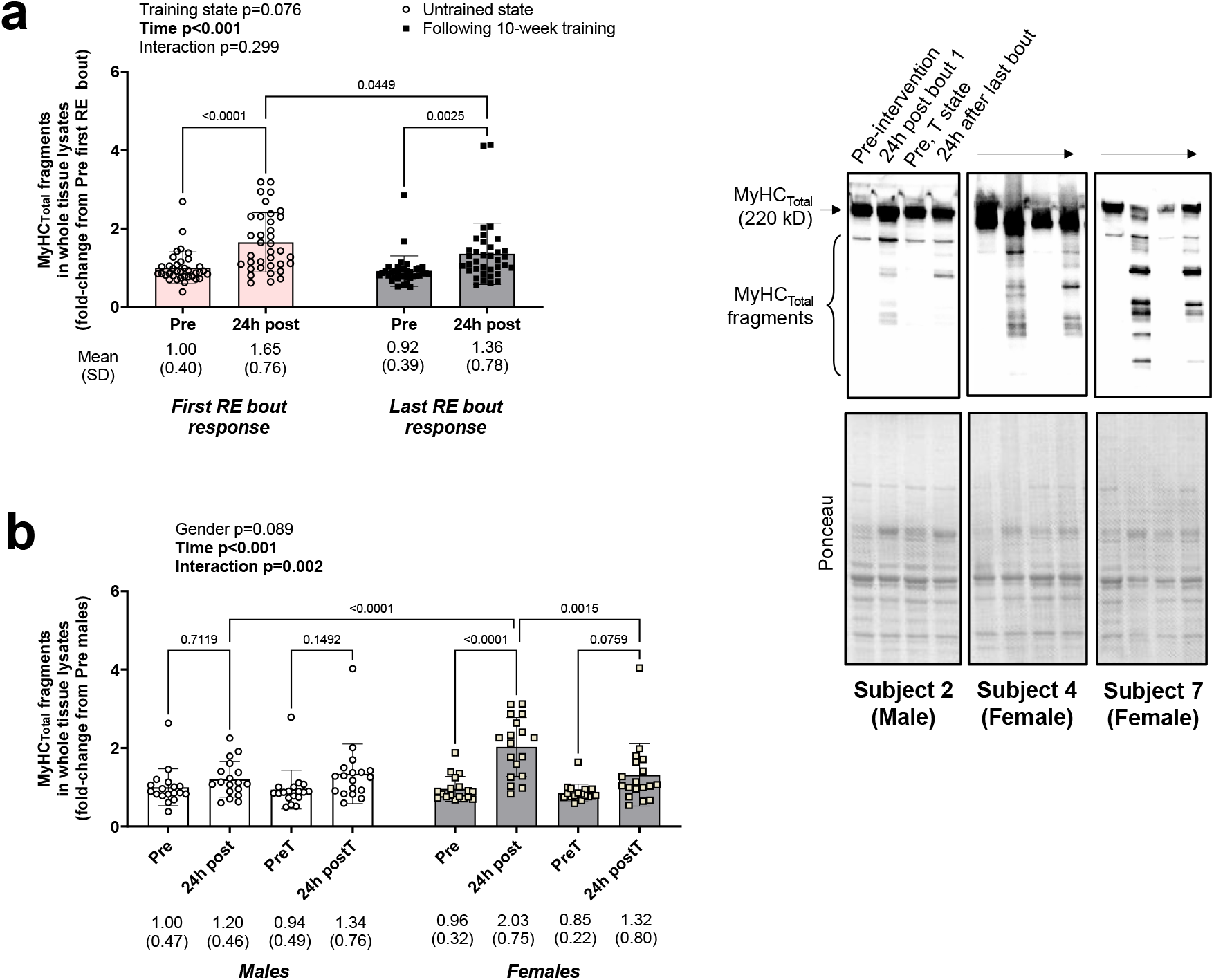
24-hour post-exercise MyHC_Total_ fragmentation in the untrained and trained states in 10weekRT participants Data from all 10weekRT participants (n=36) show that significant total myosin heavy chain (MyHC_Total_) fragmentation is evident in the whole tissue lysate 24 hours following the first/naïve resistance exercise bout (panel a). While this same 24-hour post-exercise response occurs following 10 weeks of training (24 leg extensor sessions), it is significantly attenuated. Sex analysis in 10weekRT participants (18 men and 18 women) show that the 24-hour first bout RE responses in panel a are largely driven by females (panel b). Representative immunoblots are shown for 3 participants, and data are presented as mean and standard deviation values with two-way (training state*time) ANOVA main effect and interaction p-values.

Given that there were a robust number of men and women with this study (n=18 per sex), we also examined MyHC_Total_ fragmentation responses between sexes (Figure 6b). Interestingly, a two-way (sex × time) repeated measures ANOVA indicated that significant 24-hour MyHC_Total_ fragmentation following the first/naïve bout was only evident in females (p<0.001 within and between sexes). However, this response in females was attenuated 24 hours following the last bout of training (p=0.002).

### MyHC_Total_ fragmentation is absent following a cycling bout

Figure 7 shows MyHC_Total_ fragmentation responses in the 7 participants who engaged in 60 minutes of cycling exercise. Unlike what was observed with the resistance exercise responses, there was a lack of post-exercise fragmentation 2- and 8-hours following the cycling bout.

**Figure 7.**
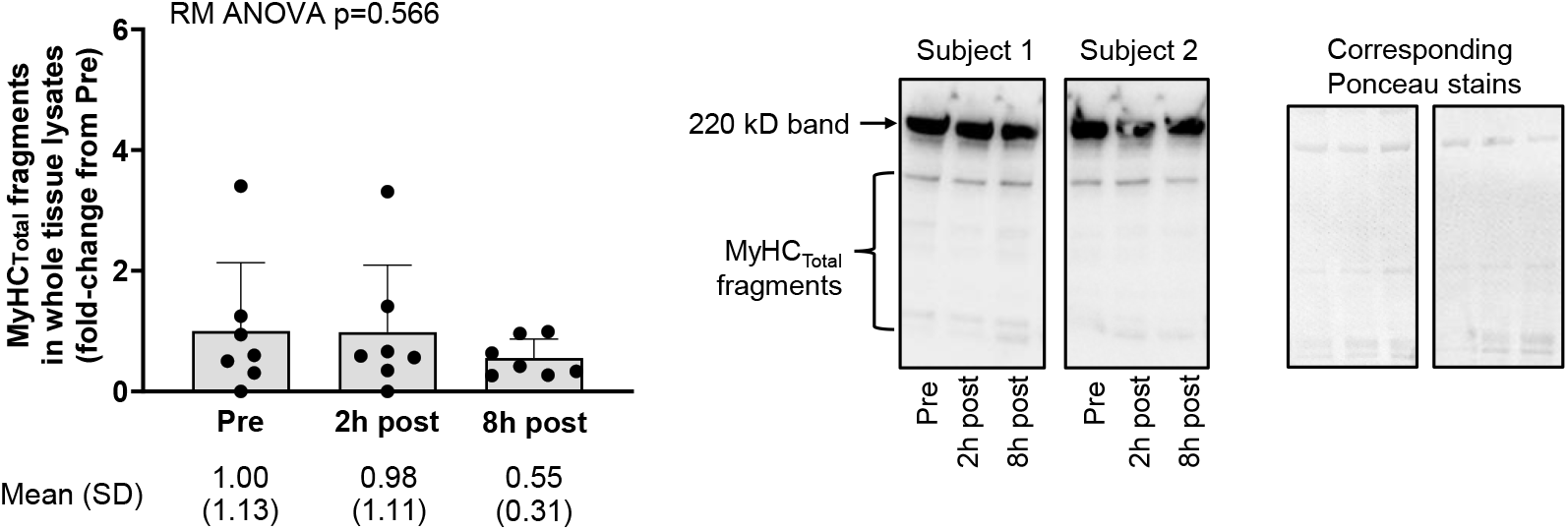
MyHC_Total_ fragmentation is absent following a cycling exercise bout Data from cycling study participants (n=7) show that total myosin heavy chain (MyHC_Total_) fragmentation is not significantly altered 2- and 8-hours following a 60-minute cycling exercise bout (panel a). Representative immunoblots are shown for 2 participants.

### MyHC_Total_ fragmentation increases following two weeks of leg immobilization

Figure 8 shows data in the 20 participants who underwent leg immobilization for two weeks. Leg immobilization led to ∼8% mid-thigh vastus lateralis atrophy (p<0.001, Figure 8a) and this coincided with a 108% increase in MyHC_Total_ fragmentation (p<0.001, Figure 8b).

**Figure 8.**
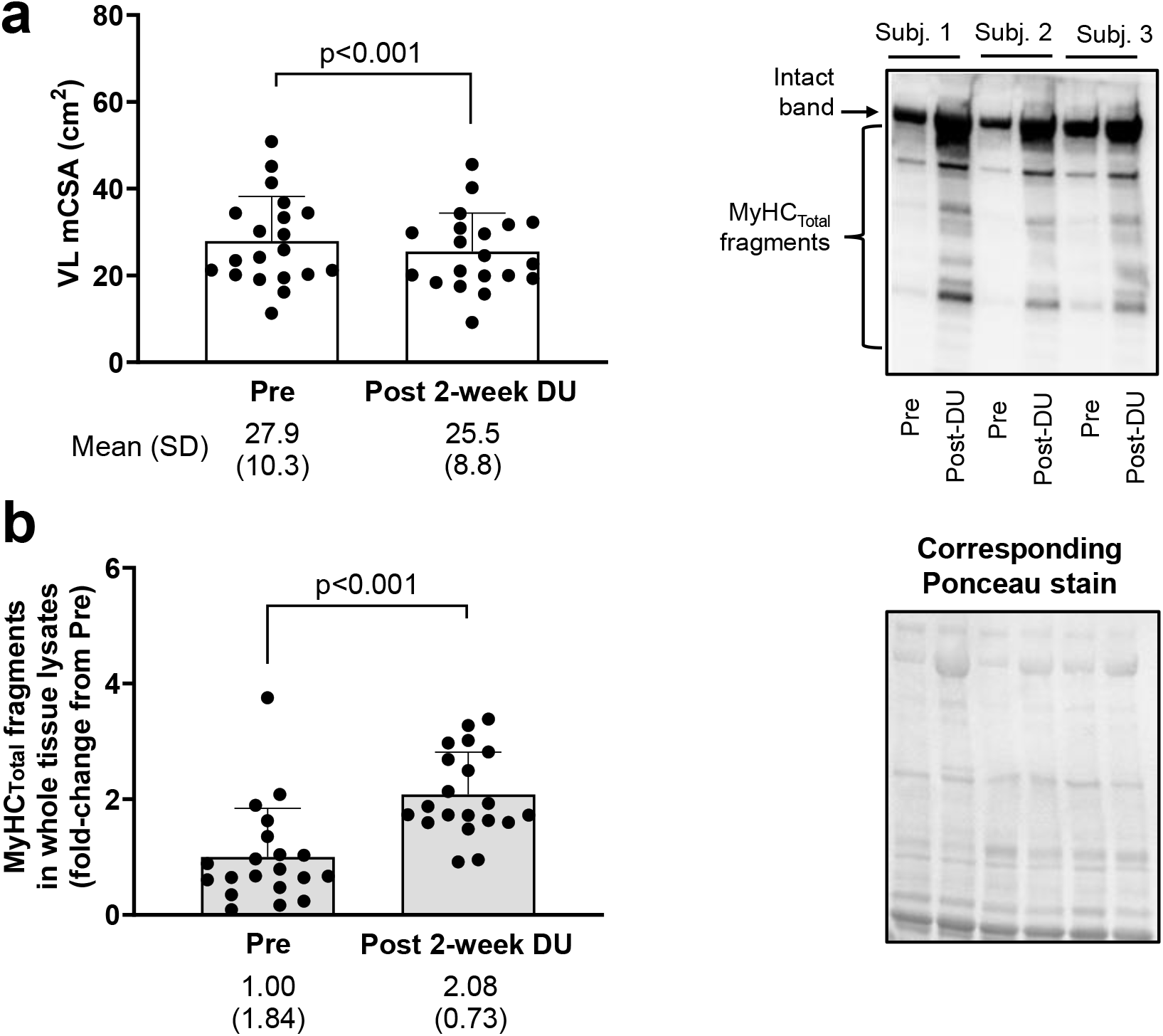
MyHC_Total_ fragmentation increases following two weeks of disuse atrophy Data from two-week disuse participants (n=20) show that VL muscle atrophy occurs with lower-limb immobilization (determined by ultrasound, panel a), and that this coincides with significant total myosin heavy chain (MyHC_Total_) fragmentation (panel b). Representative immunoblots are shown for 3 participants, and data are presented as mean and standard deviation values with dependent t-test p-values.

Although there were appreciably more men than women in this cohort (15 versus 5, respectively), statistical analyses were still performed to determine if responses were similar between sexes. Following disuse, MyHC_Total_ fragmentation increased in men (92%, p=0.002) and women (164%, p<0.001), and while the magnitude was greater in women, these responses were statistically similar between sexes (p=0.374). Likewise, no difference in VL muscle atrophy was apparent between sexes (men: -8.7±7.5%, women: -5.4±3.4%, p=0.355).

## DISCUSSION

This investigation provides evidence that RE and disuse in humans promote MyHC fragmentation. The absence of MyHC fragmentation following a cycling bout implies this process is likely a response to load-induced damage. These findings are physiologically relevant given that the MyHC protein is needed for proper muscle function and is by far the most abundant protein in skeletal muscle (i.e., ∼42% of the total muscle protein pool in college-aged men according to our recent proteomic estimates [15]). As mentioned prior, only one other study has reported that a RE bout increases titin and MyHC fragmentation in whole tissue lysates 3 hours following exercise in seven previously trained men [12]. Although the authors did not extensively pursue the significance of this finding, other studies indirectly support that the fragmentation of MyHC and other myofibrillar proteins likely occur following a RE bout. For example, Beaton et al. [16] demonstrated a loss of sarcomeric structural proteins such as desmin 4- and 24-hours following a bout of eccentric exercise in recreationally trained men. Nielson et al. [17] reported that a bout of eccentric loading leads to significant z-/m-line disruption 3-, 24- and 48-hours following exercise in untrained men; notably, z-/m-line disruption is an ultrastructural feature that may represent the release of myofilaments from intracellular structures [18]. Phillips et al. [19] reported that muscle protein breakdown (MPB) rates following RE peak at 3 hours in untrained men, and a significant elevation in MPB is also evident in these individuals 24 hours following exercise. These researchers have also reported that MPB rates are significantly elevated in resistance trained men 4 hours following a leg RE bout [20].

Ample availability of 1boutRE biopsy specimens allowed for a more expanded analysis relative to the other studies. Aside from isoform specific fragmentation patterns (discussed in the next paragraph), myofibril isolates were also analyzed for MyHC protein polyubiquitination to potentially explain the rapid disappearance of fragments by the 6-hour post-RE time point. The muscle RING finger 1 (MuRF1) and muscle atrophy F-box protein (MAFbx) E3 ligases catalyze sarcomeric protein poly-ubiquitination for degradation through the proteasome [21, 22], although there is additional evidence that polyubiquitinated protein aggregates can undergo selective lysosome degradation [23]. Hence, we hypothesized that post-RE MyHC fragments are likely polyubiquitinated. Contrary to this hypothesis, however, were the IP experiment results indicating that polyubiquitinated MyHC_Total_ fragments (normalized to the MyHC_Total_ IP signal) were not altered 3- and 6-hours post RE. Moreover, polyubiquitinated myofibril proteins were not altered at post-exercise time points and there was a lack of polyubiquitinated MyHC_Total_ fragments in the ∼50-220 kD range. Previous data published by our laboratory demonstrates that the ubiquitin antibody used in the current study readily detects proteolytic activity as observed by the accumulation of polyubiquitinated proteins in myotubes treated with a proteasome inhibitor [24]. Hence, this lends further credence that post-RE MyHC fragments are not polyubiquitinated and may be cleared from muscle through non-proteolytic mechanisms. These findings call into question how MyHC fragments are cleared from muscle. It is tempting to speculate that MyHC fragments have dissociated peptide bonds rapidly repaired and are re-associated back into myofibrils post-RE. However, enzymes facilitating this process require a catalytic domain that possesses peptide bond formation capabilities, and although non-ribosomal peptide synthetases (NRPSs) have been exist in bacteria [25], these enzymes do not exist in mammalian cells. Another explanation is that MyHC fragments are packaged into extracellular vesicles (EVs) and are released into circulation post-RE. This is not too far-fetched given that others have reported robust elevations in circulating EVs immediately post-RE [26], and MyHC has been reported to be enriched in circulating EVs [27]. However, we were not able to test this hypothesis given that blood was not obtained in 1boutRE participants. Therefore, future research is needed in further determining the fate of post-RE MyHC fragments.

The unique RE-induced MyHC I and IIa isoform fragmentation responses in the 1boutRE study participants also warrant consideration. Although the isoform responses were indeed interesting, this finding was not unanticipated given that fiber type-specific macromolecule and proteome differences have been reported (reviewed in [28]). Calpains, which are chiefly responsible for the cleavage of MyHC [29], have been reported to be differentially expressed in slow-versus fast-twitch muscle [30, 31]. Moreover, endogenous calpain inhibitors are differentially expressed in slow-versus fast-twitch muscle [32]. Hence, these fiber type differences may partially be responsible for the post-RE type I versus IIa isoform fragmentation patterns. Mechanisms aside, it is tempting to speculate how divergent post-exercise MyHC isotype fragmentation patterns associate with muscle hypertrophy outcomes. In this regard, there is generally a greater increase in type II versus type I fiber cross-sectional area in response to a variety of resistance training protocols [28, 33], and our laboratory has reported this on multiple occasions [34-37]. There is also evidence to suggest that plyometric resistance exercise leads to significantly more sarcomere damage in type II versus I fibers (∼85% versus 27%) as assessed by transmission electron microscopy [38]. This prior finding agrees in principle with observations of a more robust post-RE increase in IIa versus I isoform fragmentation. However, we temper our enthusiasm for various reasons. First, the compact pattern of IIa isoform fragmentation made it more difficult to distinguish between the intact protein and protein fragments (see lane profile in Figure 4b). Additionally, the type I isoform yielded more pre-exercise immunoreactive fragments across numerous participants compared to the type IIa isoform. Hence, the utilization of more refined approaches (e.g., longer electrophoresis run times for IIa assays) is needed to determine the extent of IIa isoform fragmentation. Future investigations parsing the mechanisms responsible for fiber type-specific differences in fragmentation and/or if these differences are associated with fiber type-specific responses to training are also warranted.

The robust MyHC_Total_ fragmentation response following two weeks of disuse atrophy in both sexes is another novel finding that is worthy of discussion. Multiple pathways catalyze MPB including calpain-mediated proteolysis, lysosome-mediated autophagy, and the ATP-dependent ubiquitin proteasome pathway [39, 40]. Although human and rodent disuse studies support an upregulation in surrogate skeletal muscle and/or blood markers related to these processes [41-46], MPB has been reported to remain unaltered or paradoxically reduced during 4 and 21 days of different disuse models in younger males [47, 48]. Indeed, this may be a limitation of tracer methods and assumptions used to assess MPB as posited by O’Reilly et al. [49]. One limitation herein is the lack of time course biopsies during leg immobilization. In this regard, an interesting interrogation would include examining MyHC fragmentation patterns one, 3, and/or 5 days following leg bracing. Notwithstanding, the current data suggest that the breakdown of MyHC occurs with VL muscle atrophy following disuse in humans, and this simple-to-execute immunoblot-based assay may serve as a viable proteolysis surrogate for future disuse studies.

Finally, the robust sample size of the 10weekRT study allowed for comparisons based on training status and sex that warrant further discussion. If indeed post-RE MyHC fragmentation is a surrogate for myofiber damage, then the Figure 6 data suggest that a naïve RE bout may elicit a greater 24-hour post-exercise damage response, as the response was attenuated in the trained state. These findings are supported by Damas et al. [50], who investigated the global transcriptome signature in nine young men. Muscle biopsies were conducted at rest and 24 hours post-resistance training (RT), both before (untrained state) and after (trained state) a 10-week resistance training program. An upregulation of genes associated with the ubiquitin-proteasome pathway (UPP), the calpain pathway, and extracellular matrix (ECM) remodeling were observed 24 hours after single RE bouts, with a more notable increase observed in the untrained state. Additionally, these results were accompanied by a reduction in muscle damage [51].

Interestingly, the attenuation of increased fragmentation in the trained state was greater for men than for women. This counters the notion that estradiol confers protection against post-exercise muscle damage, albeit considerable debate has ensued suggesting that this phenomenon is confined to rodents [52]. Moreover, sex-based differences in anabolic signaling, mRNAs associated with proteolysis, or MPS rates are minimal between young adult men and women over a 24-hour post-exercise period [53]. Women tend to experience less acute fatigue [54], and thus speculation may exist that more disruption per session limits recovery between sessions. However, while some evidence exists showing greater relative strength decrements [55, 56], a heightened post-exercise inflammatory response [57], and a heightened post-exercise blood creatine kinase response [55] compared to males, other evidence contradicts these findings [58-60]. Mechanisms potentially responsible for this sex-divergent response were not interrogated. However, the significance of this finding is questionable for a couple of reasons. First, in prior publications involving both sexes from this study demonstrated similar hypertrophic outcomes following 10 weeks of training [13, 61]. Moreover, the 24-hour post-exercise MyHC_Total_ fragmentation response to the last bout of exercise was attenuated in females and not different between sexes. Notwithstanding, the current study provides additional data to support that sex-based differences in response to RE exist and provide a further impetus to examine this area of muscle biology.

## Conclusions

In summary, MyHC fragmentation occurs in response to RE bouts and disuse atrophy in humans. A refined fragmentation response with 10 weeks of resistance training, and more refined responses in well-trained participants, suggest this an adaptive process. Importantly, we posit that this easy-to-execute immunoblot-based technique has promising utility with resistance exercise or disuse studies. More research is needed to determine how different exercise modalities (e.g., concurrent training), aging, or certain diseases that promote skeletal muscle atrophy (e.g., hyper-metabolic stress, cancer-cachexia, etc.) affect MyHC fragmentation. Research into the physiological consequences of different fragmentation responses as well as the fate of MyHC fragments is also warranted.

## ACKNOWLEGEMENTS

None of the authors disclose conflicts of interest related to these data. Funds for MyHC antibodies and associated reagents were made possible through a gift donation by Renaissance Periodization to the laboratory of M.D.R. C.A.L. was supported by The São Paulo Research Foundation (#2023/04739-2) and National Council for Scientific and Technological Development (#311387/2021-7). D.L.P. was supported by an Auburn University Presidential Research Fellowship, D.A.A. was supported aby an Auburn University Presidential Opportunity Fellowship, and M.C.M. was supported by a NIH fellowship (5T32GM141739-03). Participant compensation costs were provided as indicated in the original studies. None of the authors have financial other conflicts of interest to disclose in relation to these data. Raw data will be made available by the corresponding author upon reasonable request.

## Notes

### Competing Interest Statement

The authors have declared no competing interest.

